# The absence of yeast Pex23 and Pex29 results in a mitochondrial fusion defect

**DOI:** 10.1101/2023.12.05.570083

**Authors:** Haiqiong Chen, Rinse de Boer, Arjen M. Krikken, Fei Wu, Ida van der Klei

## Abstract

Pex23 family proteins are membrane proteins of the endoplasmic reticulum that play a role in peroxisome and lipid body formation. The yeast *Hansenula polymorpha* contains four members: Pex23, Pex24, Pex29 and Pex32. We previously showed that the loss of Pex24 or Pex32 results in severe peroxisomal defects, caused by reduced peroxisome-endoplasmic reticulum membrane contact sites. We now analyzed whether the absence of Pex23 proteins affects other organelles. Vacuoles were normal in all deletion strains. The number of lipid droplets was reduced in *pex23* and *pex29*, but not in *pex24* and *pex32*, indicating that peroxisome and lipid droplet formation require different Pex23 proteins. In *pex23* and *pex29* cells, mitochondria were fragmented and clustered. This phenotype was not suppressed by an artificial mitochondria-endoplasmic reticulum tether, indicating that the abnormalities were not caused by reduced membrane contact sites. Deletion of *DNM1* in *pex23* cells partially suppressed the phenotype. Also, the level of the mitochondrial fusion protein Fzo1 was reduced in *pex23* and *pex29* cells. These observations indicate that certain Pex23 family proteins are required for normal mitochondrial fusion.

## Introduction

Peroxisomes are organelles that contain a proteinaceous matrix and are enclosed by a single membrane. They perform several metabolic functions, depending on the organism, tissue and developmental stage. Common functions include fatty acid β-oxidation and H_2_O_2_-based respiration (Smith and Aitchison, 2013). Proteins involved in peroxisome biogenesis are called peroxins and are encoded by *PEX* genes. So far, 37 *PEX* genes have been identified (Jansen et al., 2021).

Most peroxins localize to peroxisomes. However, members of the Pex23 protein family localize to the endoplasmic reticulum (ER). Proteins of this family are characterized by a transmembrane domain and a C-terminal Dysferlin (DysF) domain (Jansen et al., 2021). The function of the DysF domain is still unknown (Bulankina and Thoms, 2020).

Pex23 family members only occur in yeast and filamentous fungi. The number of Pex23 family members varies. *Saccharomyces cerevisiae* has five (called Pex28, Pex29, Pex30, Pex31 and Pex32), while *Hansenula polymorpha* contains four Pex23 family members (Pex23, Pex24, Pex29 and Pex32) (Jansen et al., 2021).

*S. cerevisiae* Pex23 family proteins have been extensively studied. Initially, these proteins were demonstrated to play a role in de novo peroxisome biogenesis and in the formation of peroxisome-ER membrane contact sites (David et al., 2013; Mast et al., 2016). Proteomics studies showed that *S. cerevisiae* Pex29, Pex30 and Pex31 are components of a larger complex together with the ER reticulon-like proteins Rtn1, Rtn2 and Yop1 at ER-peroxisome contact sites. This protein complex defines a specialized domain of the ER where pre-peroxisomal vesicles (PPVs) bud off during de novo peroxisome formation (David et al., 2013; Farré et al., 2019). Deletion of *S. cerevisiae PEX30* or *PEX31* changes the kinetics of PPVs formation, indicating that these proteins regulate PPV formation (David et al., 2013; Joshi et al., 2016; Mast et al., 2016). ScPex30 also colocalizes with several lipid droplet (LD) biogenesis factors. Hence, the same specialized ER domain plays a role in the biogensis of PPVs and LDs (Choudhary et al., 2020; Joshi et al., 2018; Wang et al., 2018).

ScPex30 also functions at nuclear–vacuolar junctions (NVJs). To fulfil its different functions ScPex30 forms complexes with other members of the Pex23 protein family. For the formation of peroxisome-ER contact sites, Pex30 associates with Pex28 and Pex32, while it associates with Pex29 in NVJs (Ferreira and Carvalho, 2021).

We previously studied the four members of the Pex23 family of the yeast *H. polymorpha.* We showed that HpPex24 and HpPex32 play key roles in peroxisome biogenesis and required for the formation of ER-peroxisome contacts (Wu et al., 2020). The latter is underscored by the observation that the peroxisomal defects of *pex24* and *pex32* mutants could be suppressed by the introduction of an artificial ER-peroxisome tethering protein. Interestingly, cells lacking HpPex29 showed no peroxisome phenotype, whereas the absence of HpPex23 has only a minor effect on peroxisomes (Wu et al., 2020).

While HpPex24 and HpPex32 accumulate at peroxisome-ER contact sites, HpPex23 and HpPex29 localize to multiple regions of the ER. Moreover, like ScPex30, HpPex23 also accumulates at NVJs (Wu et al., 2020).

Our observation that cells lacking HpPex29 do not show any peroxisomal defect, suggests that Pex23 family proteins may have additional functions. In the current study we show that the absence of HpPex23 and HpPex29 affects LD formation and mitochondria fusion. Importantly, in *H. polymorpha* defects in peroxisome and LD biogenesis are not linked, because HpPex29 is important for LD biogenesis, but does not play a role in peroxisome biogenesis.

## RESULTS

### Deletion of *PEX23* or *PEX29* affects mitochondria and LDs

To study whether the absence of *H. polymorpha* Pex23 family proteins affects other organelles in addition to peroxisomes, we analysed the morphology of vacuoles, LDs and mitochondria in *pex23*, *pex24*, *pex29* and *pex32* cells. Fluorescence microscopy (FM) revealed that in all deletion strains vacuole morphology (marked with FM4-64) was similar as in the WT control (Fig. 1A). In contrast, upon staining the cells with BODIPY 493/503 LDs were easily detected in WT, *pex24* and *pex29* cells, whereas in *pex23* and *pex29* cells only a few faint spots could be detected. The latter may reflect decreased LD numbers or changes in their lipid composition, which could influence BODIPY 493/505 staining. We therefore also used Erg6-GFP as a protein marker for LDs. In cells producing endogenous Erg6-GFP the number of fluorescent spots was reduced in *pex23* and *pex29* relative to *pex24, pex32* and WT controls (Fig. 1A, B). Western blotting showed that Erg6-GFP protein levels were similar in all five strains tested, indicating that the lower number of LDs detected in *pex23* and *pex29* is not due to reduced levels of the marker Erg6-GFP. We therefore conclude that the decrease in BODIPY 493/505 or Erg6-GFP marked spots is indeed reflecting reduced LD abundance in *pex23* and *pex29* cells (Fig. 1C).

**Figure 1.**
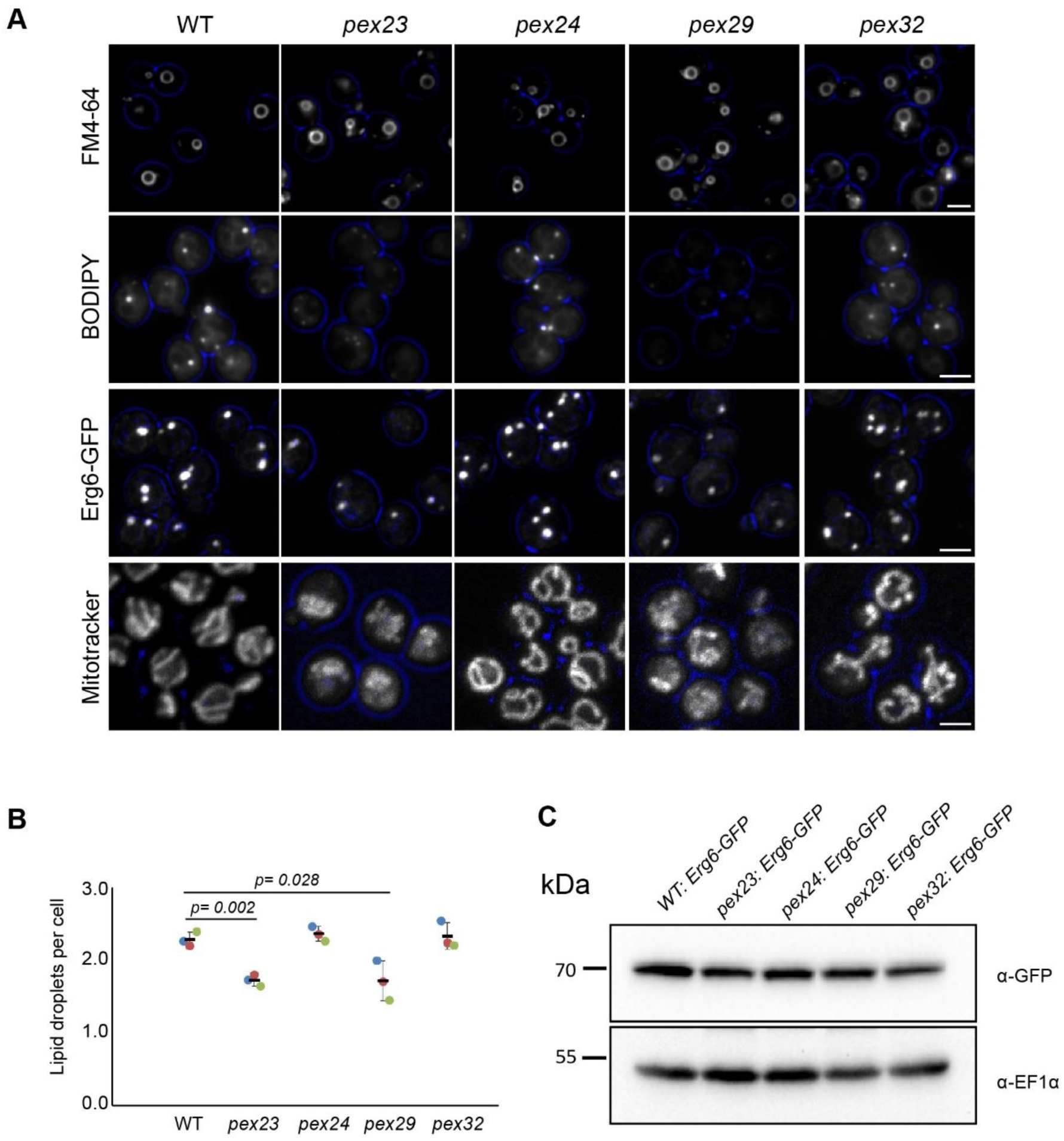
Deletion of *PEX23* or *PEX29* alters LDs and mitochondria. (A) FM images (vacuoles) and Confocal Laser Scanning Microscopy (CLSM) images (LDs and mitochondria) of glucose-grown cells of the indicated deletion strains. Cells were stained with FM4-64 to visualize vacuoles, BODIPY 493/503 for LDs or Mitotracker Orange for mitochondria. In addition, cells that produce Erg6-GFP were analysed to mark LDs. Scale bar: 2 µm. (B) Quantification of LD numbers based on the number of Erg6-GFP puncta in Z-stack confocal images. (*n*=3 using 900 cells from three independent experiments) (C) Western blot analysis of the Erg6-GFP in the indicated strains. Blots were decorated with anti-GFP or anti-EF1α. EF1α was used as a loading control.

Interestingly, *pex23* and *pex29* deletion cells also displayed aberrant mitochondrial profiles, based on FM analysis of Mitotracker-stained cells. In *pex24* and *pex32* cells, which show severe peroxisomal defects (Wu et al., 2020), mitochondrial morphology was similar to the WT control (Fig. 1A).

Summarizing, our FM data show that the absence of Pex23 and Pex29, but not Pex24 or Pex29, affects mitochondria and LDs.

Aberrant LD formation was previously reported for *S. cerevisiae* cells lacking Pex30, a protein that also is a member of the Pex23 family (Choudhary et al., 2020; Joshi et al., 2018; Wang et al., 2018). We now for the first time observed alterations in mitochondrial morphology in yeast cells lacking specific Pex23 family members.

### Fragmentation and clustering of mitochondria in *pex23* and *pex29* cells

To better understand the alterations in mitochondria morphology, electron microscopy (EM) analysis was performed. As shown in Fig. 2A, mitochondrial profiles were present throughout the cells in WT, *pex24* and *pex32* cells, in line with the FM observations. Quantification of the number of mitochondrial profiles per cell section revealed that thin sections of WT cells contain up to 4 mitochondrial profiles, with most sections containing 1 to 3 mitochondrial profiles. A very similar distribution was observed in sections of *pex24* cells. However, in sections of *pex23* and *pex29* cells, and to a lesser extend of *pex32* cells, a higher number of mitochondrial profiles was observed (Fig. 2A).

**Figure 2.**
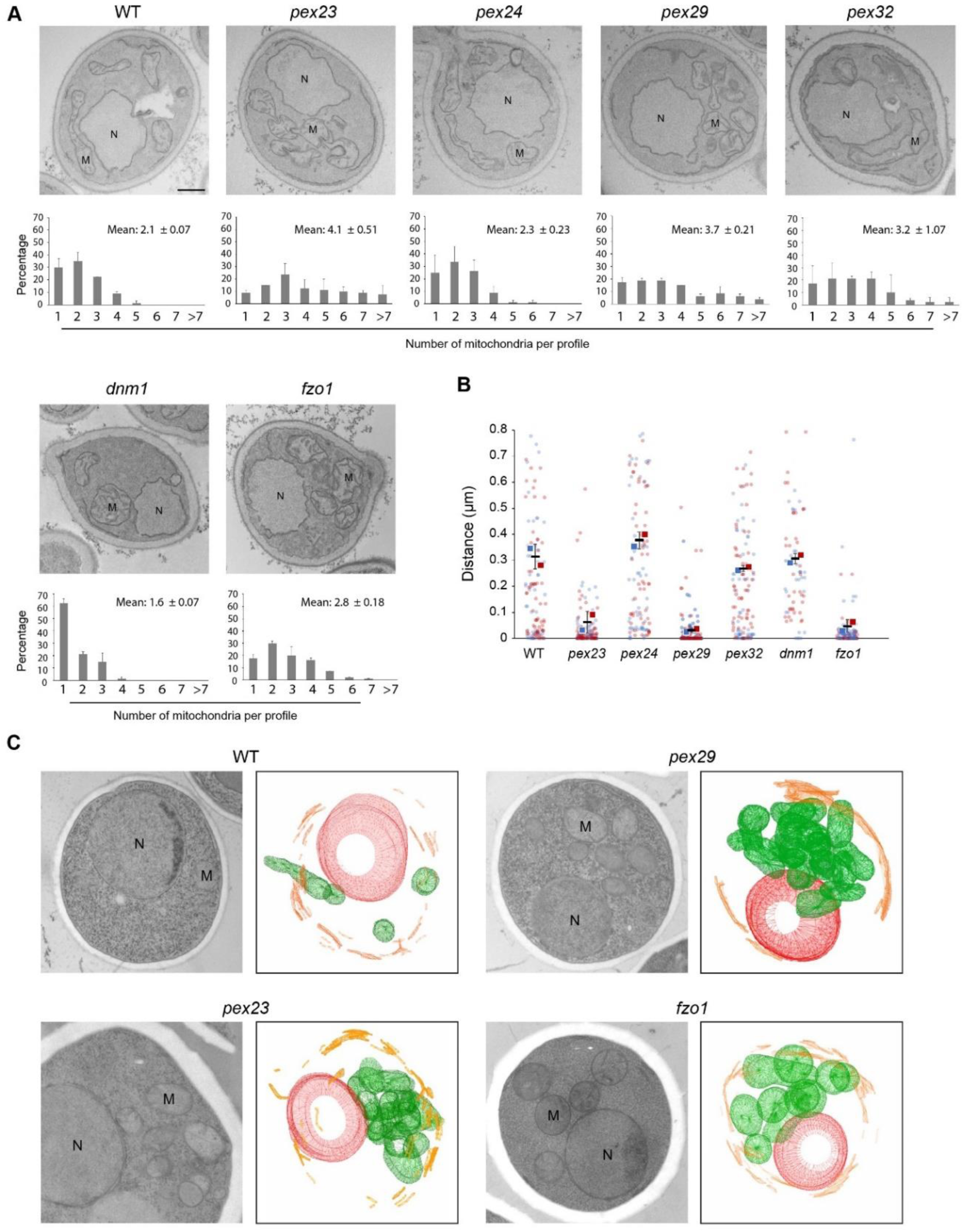
*H. polymorpha pex23* and *pex29* cells contain a cluster of multiple mitochondria. (A) EM images of thin sections of KMnO_4_-fixed glucose-grown cells of the indicated strains, and the number of mitochondria per profile have been quantified. Data are mean ± SD of two independent experiments (*n*=2 using 40 random sections from each experiment). M, mitochondrion; N, nucleus. Scale bar: 500 nm. (B) Quantification of the distance between mitochondria based on EM images of indicated strains. (C) 3D reconstructions of serial sections of cryo-fixed cells. See supplemental material for the raw data. M - mitochondria (green), N – nucleus (red), ER- endoplasmic reticulum (orange).

Subsequently, we quantified the distance between mitochondrial profiles, which revealed that mitochondria are more clustered in *pex23* and *pex29* mutants relative to *pex24*, *pex32* and WT cells (Fig. 2B). EM analysis of serial sections of *pex23* and *pex29* cells confirmed that these cells contain multiple mitochondria, which are highly clustered (Fig. 2C, Fig. S1).

Yeast Dnm1 and Fzo1 are crucial proteins in mitochondrial fission and fusion, respectively (Hermann et al., 1998; Sesaki and Jensen, 1999). As expected, the number of mitochondrial profiles in thin sections of *H. polymorpha dnm1* cells is strongly reduced (Fig. 2A), whereas an increase is observed in *fzo1* cell sections. In *dnm1* cells, the distance between mitochondria is similar as in the WT control. However, in *fzo1* cells mitochondria are much more clustered, similar as observed in *pex23* and *pex29* cells (Fig. 2B).

In conclusion, the mitochondrial phenotype of *pex23* and *pex29* cells is comparable with that of *fzo1* cells and characterized by the presence of more mitochondria which are clustered.

### Mitochondrial activity is reduced in *pex23* and *pex29* cells

To determine whether the altered mitochondrial morphology results in decreased mitochondrial activity, we stained glucose-grown cells with Rhodamine123 (Rh123). Rh123 is a cell-permeable cationic green fluorescent dye that is taken up by active mitochondria and commonly used for the detection of the mitochondrial membrane potential (Drakulic et al., 2005; Ludovico et al., 2001). As shown in Fig. 3A, the Rh123 fluorescence intensities are reduced in *pex23* and *pex29* cells compared to *pex24*, *pex32* and WT control cells (Fig.3B). Moreover, the absence of Pex23 and Pex29, but not of Pex24 and Pex32, resulted in slightly retarded of growth (Fig.3B).

**Figure 3.**
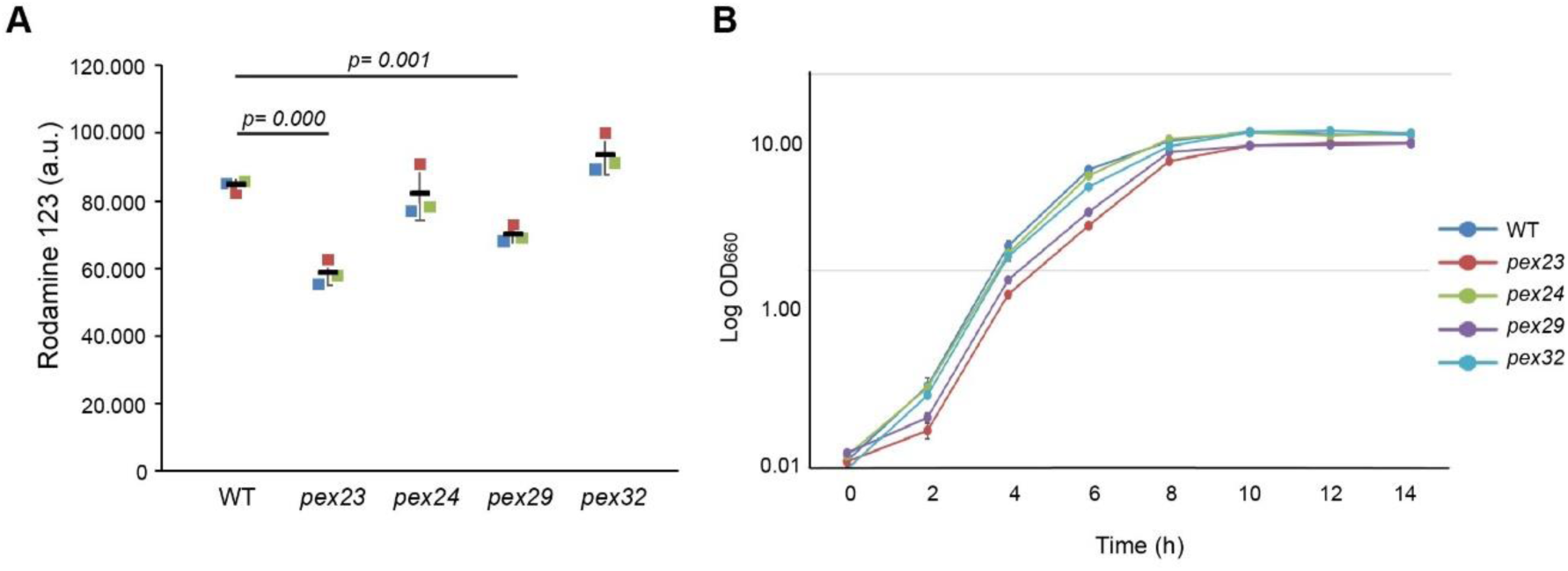
The absence of Pex23 and Pex29 results in reduced mitochondrial activities. (A) Fluorescence intensities of glucose-grown cells stained with Rh123 and measured by flow cytometry. Data are presented from three independent experiments. (B) Growth curves of cells in glucose medium. The error bars represent SD from three independent experiments.

### Enhanced ER-mitochondria contacts do not suppress the mitochondrial phenotype of *pex23* and *pex29* cells

All *H. polymorpha* Pex23 family members localize to the ER. Pex24 and Pex32 are predominantly present at peroxisome-ER contact sites, but Pex23 also accumulates at nucleus–vacuole junctions (NVJs). These observations suggest that members of the Pex23 protein family may be common ER-localized contact site resident proteins (Wu et al., 2020).

Contact sites between the ER and mitochondria regulate mitochondrial fission and fusion in *S. cerevisiae* (Abrisch et al., 2020; Prinz et al., 2019). We therefore analyzed whether Pex23 also accumulated at ER-mitochondria contact sites using correlative light and electron microscopy (CLEM). Pex23-GFP was slightly overproduced to obtain sufficient fluorescence signal to enable CLEM analysis. As shown in Fig. 4A, Pex23-GFP especially occurred at regions where peripheral ER is present (Fig. 4A). Mitochondria also form contacts with peripheral ER, hence Pex23 is present at mitochondria-ER contacts. However, Pex23-GFP does not accumulate at ER-mitochondria contacts. In line with our earlier report, Pex23-GFP also accumulates at NVJs (Fig. 4A).

**Figure 4.**
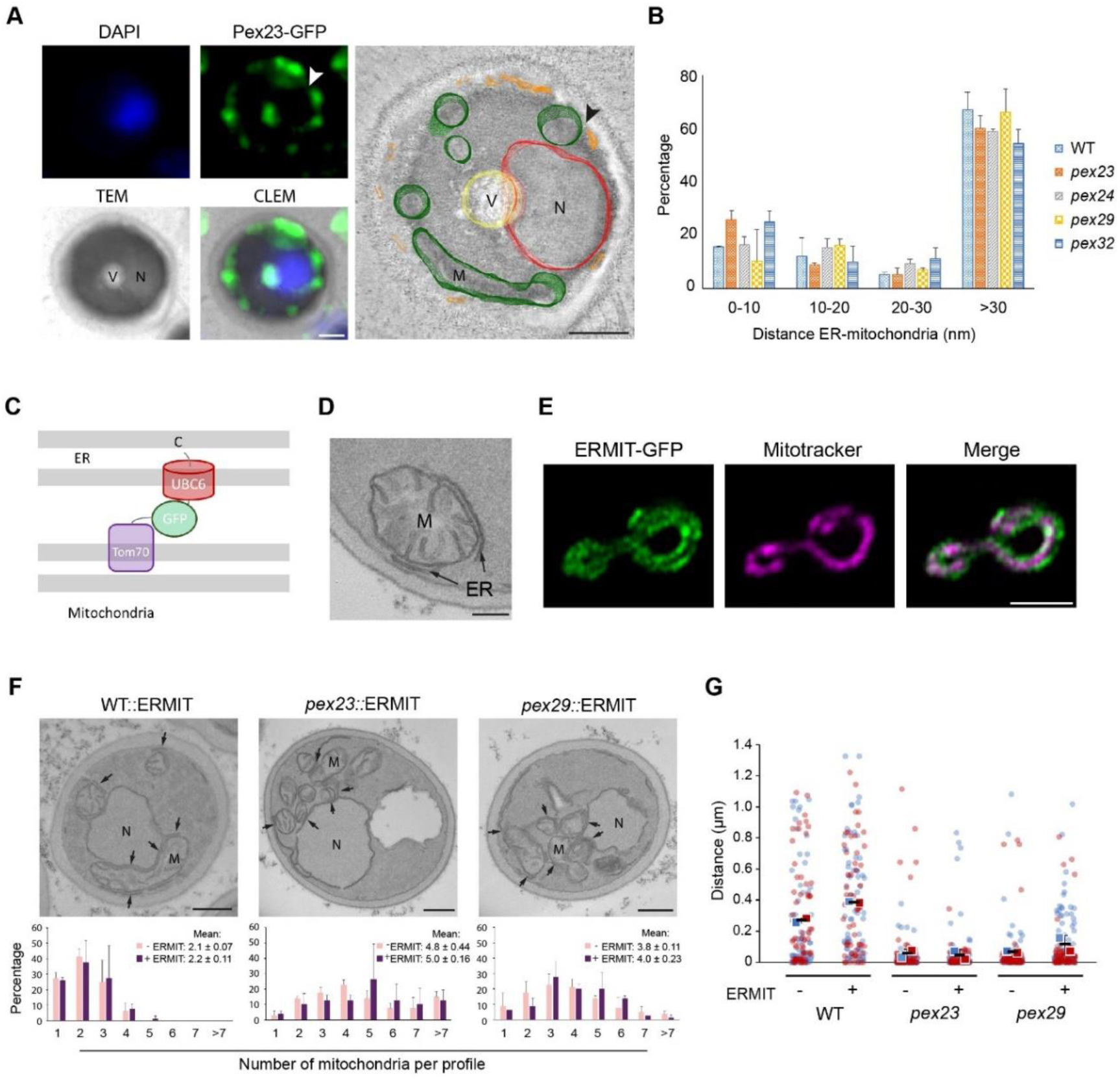
ERMIT-GFP does not suppress the mitochondrial phenotypes of *pex23* and *pex29* cells. (A) CLEM of cells producing Pex23-GFP under control of the P*_AOX_*. The nucleus is stained with 4′,6-diamidino-2-phenylindole (DAPI). The arrowhead indicates the most intense GFP spot in the GFP image and the corresponding position in the rendered volume. Cells were induced for 1.5 h on glycerol medium. V - vacuole, N - nucleus, M - mitochondrion. (B) Quantification of the distance between the ER and mitochondrial membranes in the indicated strains using sections of KMnO_4_-fixed cells. (C) Schematic representation of the artificial tether protein ERMIT. (D) EM image of a thin section of a KMnO_4_-fixed glucose-grown WT cell producing ERMIT. (E) Single plane CLSM Airyscan image of WT cell producing the ERMIT tether. Mitochondria are stained with Mitotracker Orange. (F) EM images of thin sections of KMnO_4_-fixed glucose-grown cells of the indicated strains, and the number of mitochondrial profiles per section. Data are mean ± SD of two independent experiments (n=2 using 40 random sections from each experiment). M, mitochondrion; N, nucleus. Scale bar: 500 nm. (G) Quantification of the distance between mitochondria in indicated cells.

EM analysis of thin sections revealed that the percentage of mitochondria that are closely associated with the ER is similar in all four mutant strains and the WT control (Fig. 4B). This indicates that in the absence of Pex23 family proteins physical ER-mitochondrial contacts are not reduced.

As a complementary approach, we tested the effect of introducing an artificial ER-mitochondrion tether (called ERMIT). Previous studies in *S. cerevisiae* revealed that an artificial tether can suppress the phenotype of mutants lacking a component of the ERMES complex, which is important for ER-mitochondria contacts (Kornmann et al., 2009). ERMIT consists of full length Tom70 at the N-terminus, followed by GFP and the C-terminal domain of the ER tail anchored protein Ubc6 (Fig. 4C). EM analysis showed that extensive ER-mitochondria contacts and contacts between the nuclear envelope and mitochondria occurred in cells producing this artificial tether (Fig. 4D). FM analysis showed that ERMIT fully co-localized with mitochondria (Fig. 4E).

Quantification of mitochondrial profiles on EM sections showed no change upon introduction of ERMIT (Fig. 4F). Also, the distances between mitochondria did not increase in cells of any of the four deletion strains. Hence the level of clustering was unchanged upon introduction of ERMIT (Fig. 4G). A slight increase in distance between mitochondria was observed for the WT control (T test p = 0.01) (Fig. 4G). Possibly, the artificial ER-mitochondrial contacts keep the mitochondria a bit more apart from each other.

Summarizing our results show that the mitochondrial phenotypes of *pex23* and *pex29* are unlikely due to a change in physical contacts between the ER and mitochondria.

### Mitochondrial fission and/or fusion are altered in *pex23* cells

The increased number of mitochondrial profiles in *pex23* and *pex29* cells may be due to decreased mitochondrial fusion or enhanced organelle fission. Western blot analysis of cells producing Dnm1-GFP or Fzo1-GFP under control of their endogenous promoters, revealed that the levels of Fzo1-GFP were reduced in *pex23* and *pex29* cells relative to the WT control. The levels of Dnm1-GFP were only reduced in *pex32* cells. FM analysis showed that in *pex23* cells the localisation of Fzo1 and Dnm1 had not changed.

Based on these observations we conclude that mitochondrial fusion is reduced in *pex23*, *pex29* and *pex32* cells due to reduced levels of Fzo1. Moreover, most likely mitochondrial fission is reduced in *pex32* cells due to lower Dnm1 levels.

In *S. cerevisiae*, deletion of *DNM1* in *fzo1* cells restored the mitochondria phenotype (Sesaki and Jensen, 1999). We reasoned that if a similar suppression by *DNM1* deletion also occurs in *pex23* cells, this would support a direct or indirect role of Pex23 in mitochondrial fusion. To generate a *dnm1 pex23* double mutant, we deleted *DNM1* in *pex23*. Quantification of mitochondrial profiles in EM sections showed that the number of mitochondrial profiles decreased again (Fig. 5A). In addition, the distance between the mitochondria profiles decreased, indicative for less clustering (Fig. 5B). Also, mitochondrial activity increased and cell growth on glucose was restored in the double deletion strain (Fig. 5C). Taken together these data show that deletion of *DNM1* in *pex23* largely suppressed the mitochondrial phenotype of *pex23* cells.

**Figure 5.**
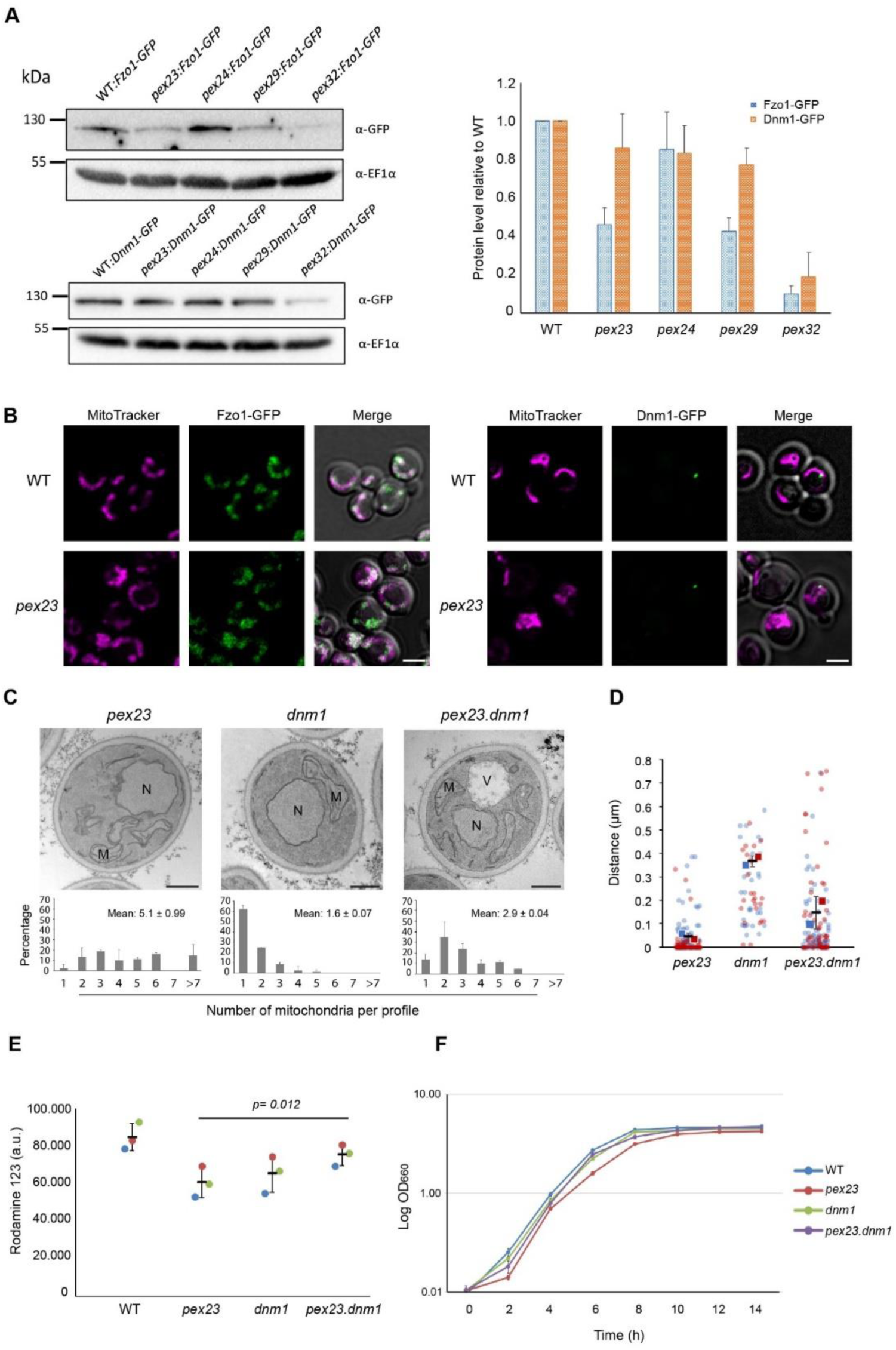
The absence of Pex23 affects mitochondrial fusion. (A)Western blot analysis and quantification of the indicated proteins in glucose-grown cells that produce endogenous Fzo1-GFP or Dnm1-GFP. Blots were decorated with anti-GFP or anti-EF1α antibodies. EF1α was used as a loading control. The protein levels of WT cells were set to 1. The error bars represent SD from three independent experiments. (B) Confocal (Airyscan, single plane) images showing the localization of Fzo1-GFP and Dnm1-GFP in glucose-grown WT and *pex23* cells. Mitochondria are marked with Mitotracker Red. Scale bar: 2 µm. (C) EM images of thin sections of KMnO_4_-fixed glucose-grown cells of the indicated strains, and the number of mitochondrial profiles per cell section. Data are mean ± SD of two independent experiments (n=2 using 40 random sections from each experiment). M, mitochondrion; N, nucleus. Scale bar: 500 nm. (D) Quantification of the distance between mitochondria in indicated strains. Data are mean ± SD of two independent experiments (n=2 using 40 random sections from each experiment). (E) Fluorescence intensities of glucose-grown cells stained with Rh123 were measured by flow cytometry. The presented data are from three independent experiments. The error bars represent SD from three independent experiments. (F) Growth curves of the indicated strains in mineral media containing glucose. The error bars represent SD from three independent experiments.

Taken together these results show that in *pex23* cells Fzo1 levels and mitochondrial fusion is reduced. Like in *pex23* cells, the Fzo1 protein levels are also reduced in *pex29* cells, hence the phenotype of this mutant most likely is also related to a mitochondrial fusion defect.

## Discussion

In this study, we show for the first time that the absence of members of the Pex23 protein family can lead to alterations in mitochondrial morphology and activity. Our data indicate that deletion of *PEX23* or *PEX29* in the yeast *H. polymorpha* leads to aberrant mitochondrial morphology. In the same two mutants the number of LDs is also reduced, which is in line with previous findings in *S. cerevisiae pex30* cells, which contain less and smaller LDs (Joshi et al., 2018; Wang et al., 2018).

However, different from what has been reported for *S. cerevisiae*, peroxisome biogenesis and LD formation appeared not to be linked in *H. polymorpha*. In *S. cerevisiae* the same, specialized subdomain of the ER, which contains Pex30 and proteins involved in LD formation, is involved in peroxisome and LD formation (Choudhary et al., 2020; Joshi et al., 2018; Wang et al., 2018). However, we here show that in *H. polymorpha* Pex24 and Pex32 have crucial roles in peroxisome biology (Wu et al., 2020), while Pex23 and Pex29 are important for LD formation (this study). Moreover, these Pex23 family proteins localize to different regions of the ER, namely at peroxisome-ER contacts (HpPex24 and HpPex32), or other regions of the ER, including NVJs (HpPex23, HpPex29) (Wu et al., 2020).

The altered mitochondrial morphology in *H. polymorpha pex23* and *pex29* cells is not indirectly due to peroxisome biogenesis defects, because *pex24* cells, which show very severe peroxisome biogenesis defects, did not exhibit mitochondrial abnormalities. Conversely, in *pex29* cells, which show mitochondrial defects, peroxisome biogenesis is normal (Wu et al., 2020).

EM analysis of *pex23* and *pex29* cells revealed an increased number of mitochondria that are highly clustered. This phenotype is most likely due to reduced mitochondrial fusion. In *S. cerevisiae* a block in mitochondrial fusion, caused by the absence of Fzo1, results in fragmented mitochondrial structures that form a cluster in one area of the cell (Hermann et al., 1998). We observed a similar phenotype for *H. polymorpha fzo1* (Fig. 2A), which highly resembled the phenotype of *H. polymorpha pex23* and *pex29* cells.

Deletion of *DNM1* in *H. polymorpha pex23* cells reduced the mitochondrial phenotype (i.e. less mitochondria and less clustering). This further points to a reduction of mitochondrial fusion, because deletion of *DNM1* in *S. cerevisiae* fzo*1* partially suppressed the phenotype (Yang et al., 2021).

The fusion defect most likely is caused by reduced Fzo1 levels in *H. polymorpha pex23* and *pex29* cells (Fig. 5A). The levels of Dnm1, responsible for mitochondrial fission, were normal in these strains. Why Fzo1 levels are reduced remains unknown. In *S. cerevisiae* Fzo1 levels are highly regulated by and include highly regulated degradation by the ubiqitin proteasome system (Cavellini et al., 2017).

Cells of the *PEX32* deletion strain also showed increased mitochondrial numbers, however these organelles were not clustered. In these cells both Fzo1 and Dnm1 levels are reduced, most likely resulting in less fision and fusion of the organelles. Why the the levels of these proteins are reduced in this mutant is still unknown.

In conclusion, we show that proteins of the Pex23 family proteins play a role not only in peroxisome and LD formation, but also in mitochondrial dynamics. This highlights the importance and functional diversity of this protein family.

## Materials and methods

### Strains and growth conditions

*H. polymorpha* cells were grown in batch cultures at 37 ℃ on mineral medium (Van Dijken et al., 1976) supplemented with 0.5% glucose or 0.25% glycerol as carbon source. When required, leucine was added to a final concentration of 60 μg/mL. For growth on plates, Yeast extract–Peptone–Dextrose (YPD) media (1% yeast extract, 1% peptone and 1% glucose) were supplemented with 2% agar. Transformants were selected using 100 μg/mL zeocin (Invitrogen), or 100 μg/mL nourseothricin (WERNER BioAgents) or 300 μg/mL hygromycin (Invitrogen).

The *Escherichia coli* strain DH5α used for cloning. *E. coli* cells were grown at 37 ℃ in Luria broth (LB) media (1% Bactotryptone, 0.5% Yeast Extract and 0.5% NaCl) supplemented with 100 μg/mL ampicillin or 50 µg/mL kanamycin. For plates 2% agar was added.

### Construction of *H. polymorpha* strains

The strains, plasmids and primers used in this study are listed in Table 1, 2 and 3 respectively. Plasmids integration was performed as described previously (Faber et al., 1994). All integrations were confirmed by PCR (Thermo Fisher Scientific). Gene deletions were confirmed by PCR and Southern blotting.

**Table 1.** Strains used in this study.

| Strain | Characteristics | Reference |
| --- | --- | --- |
| WT (*yku80*) | NCYC495 *yku80* deletion strain; *leu 1.1*, | (Saraya et al., 2012) |
| *pex23* | *yku80* with *PEX23* deletion strain; *leu 1.1*, Zeo<sup>R</sup> | (Wu et al., 2020) |
| *pex24* | *yku80* with *PEX24* deletion strain; *leu 1.1*, Zeo<sup>R</sup> | (Wu et al., 2020) |
| *pex29* | *yku80* with *PEX29* deletion strain; *leu 1.1*, Zeo<sup>R</sup> | (Wu et al., 2020) |
| *pex32* | *yku80* with *PEX32* deletion strain; *leu 1.1*, Zeo<sup>R</sup> | (Wu et al., 2020) |
| *dnm1* | *yku80* with *DNM1* deletion strain; *leu 1.1*, Hph<sup>R</sup> | (Manivannan et al., 2013) |
| *fzo1* | *yku80* with *DNM1* deletion strain; *leu 1.1*, Nat<sup>R</sup> | This study |
| WT::Pex23-GFP | *yku80* with pHIPZ-*PEX23*-mGFP; *leu 1.1*, *URA3*, Zeo<sup>R</sup> | (Wu et al., 2020) |
| WT: P<sub>AOX</sub>Pex23-GFP::DsRed-SKL | Pex23-mGFP with pHIPN4-DsRed-SKL; *leu 1.1*, *URA3*, Zeo<sup>R</sup>, Nat<sup>R</sup> | This study |
| WT: Erg6-mGFP | *yku80* with pHIPN-Erg6-mGFP; *leu 1.1*, Nat<sup>R</sup> | This study |
| *pex23*: Erg6-mGFP | *pex23* with pHIPN-Erg6-mGFP; *leu 1.1*, Nat<sup>R</sup> | This study |
| *pex24*: Erg6-mGFP | *pex24* with pHIPN-Erg6-mGFP; *leu 1.1*, Nat<sup>R</sup> | This study |
| *pex29*: Erg6-mGFP | *pex29* with pHIPN-Erg6-mGFP; *leu 1.1*, Nat<sup>R</sup> | This study |
| *pex32*: Erg6-mGFP | *pex32* with pHIPN-Erg6-mGFP; *leu 1.1*, Nat<sup>R</sup> | This study |
| WT: ERMIT | *yku80* with pHIPN18-Tom70-mGFP-Ubc6; *leu 1.1*, Nat<sup>R</sup> | This study |
| *pex23*: ERMIT | *pex23* with pHIPN18-Tom70-mGFP-Ubc6; *leu 1.1*, Nat<sup>R</sup> | This study |
| *pex29*: ERMIT | *pex29* with pHIPN18-Tom70-mGFP-Ubc6; *leu 1.1*, NAT<sup>R</sup> | This study |
| *pex23 dnm1* | *pex23* with integration of *DNM1* deletion cassette; Zeo<sup>R</sup> | This study |
| WT: Fzo1-GFP | *yku80* with pHIPN-Fzo1-mGFP; *leu 1.1*, Nat<sup>R</sup> | This study |
| *pex23*: Fzo1-GFP | *pex23* with pHIPN-Fzo1-mGFP; *leu 1.1*, Nat<sup>R</sup> | This study |
| *pex24*: Fzo1-GFP | *pex24* with pHIPN-Fzo1-mGFP; *leu 1.1*, Nat<sup>R</sup> | This study |
| *pex29*: Fzo1-GFP | *pex29* with pHIPN-Fzo1-mGFP; *leu 1.1*, Nat<sup>R</sup> | This study |
| *pex32*: Fzo1-GFP | *pex32* with pHIPN-Fzo1-mGFP; *leu 1.1*, Nat<sup>R</sup> | This study |
| WT: Dnm1-GFP | *yku80* with pHIPN-Dnm1-mGFP; *leu 1.1*, Nat<sup>R</sup> | This study |
| *pex23*: Dnm1-GFP | *pex23* with pHIPN-Dnm1-mGFP; *leu 1.1*, Nat<sup>R</sup> | This study |
| *pex24*: Dnm1-GFP | *pex24* with pHIPN-Dnm1-mGFP; *leu 1.1*, Nat<sup>R</sup> | This study |
| *pex29*: Dnm1-GFP | *pex29* with pHIPN-Dnm1-mGFP; *leu 1.1*, Nat<sup>R</sup> | This study |
| *pex32*: Dnm1-GFP | *pex32* with pHIPN-Dnm1-mGFP; *leu 1.1*, Nat<sup>R</sup> | This study |

**Table 2.**
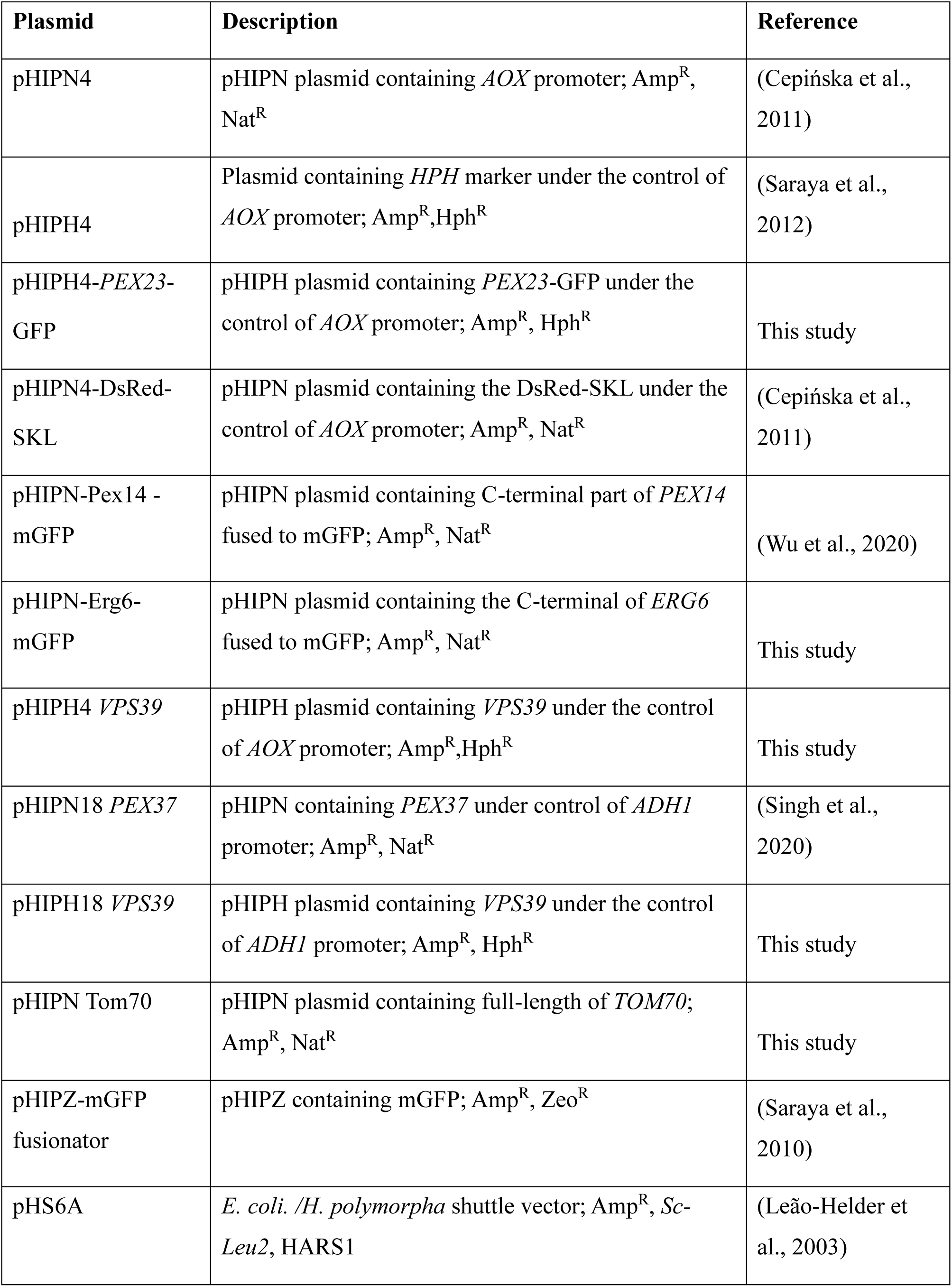
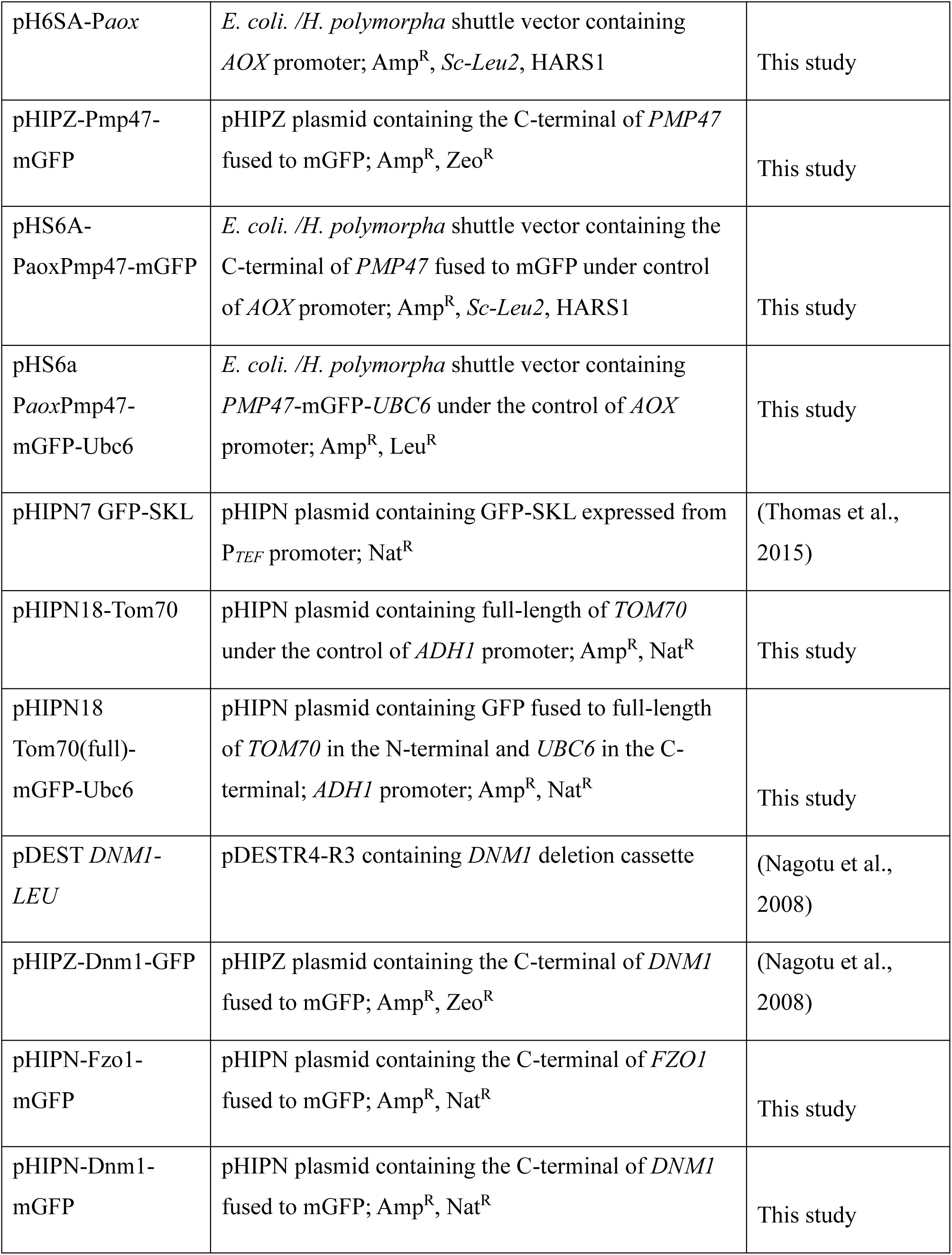
Plasmids used in this study.

**Table 3.**
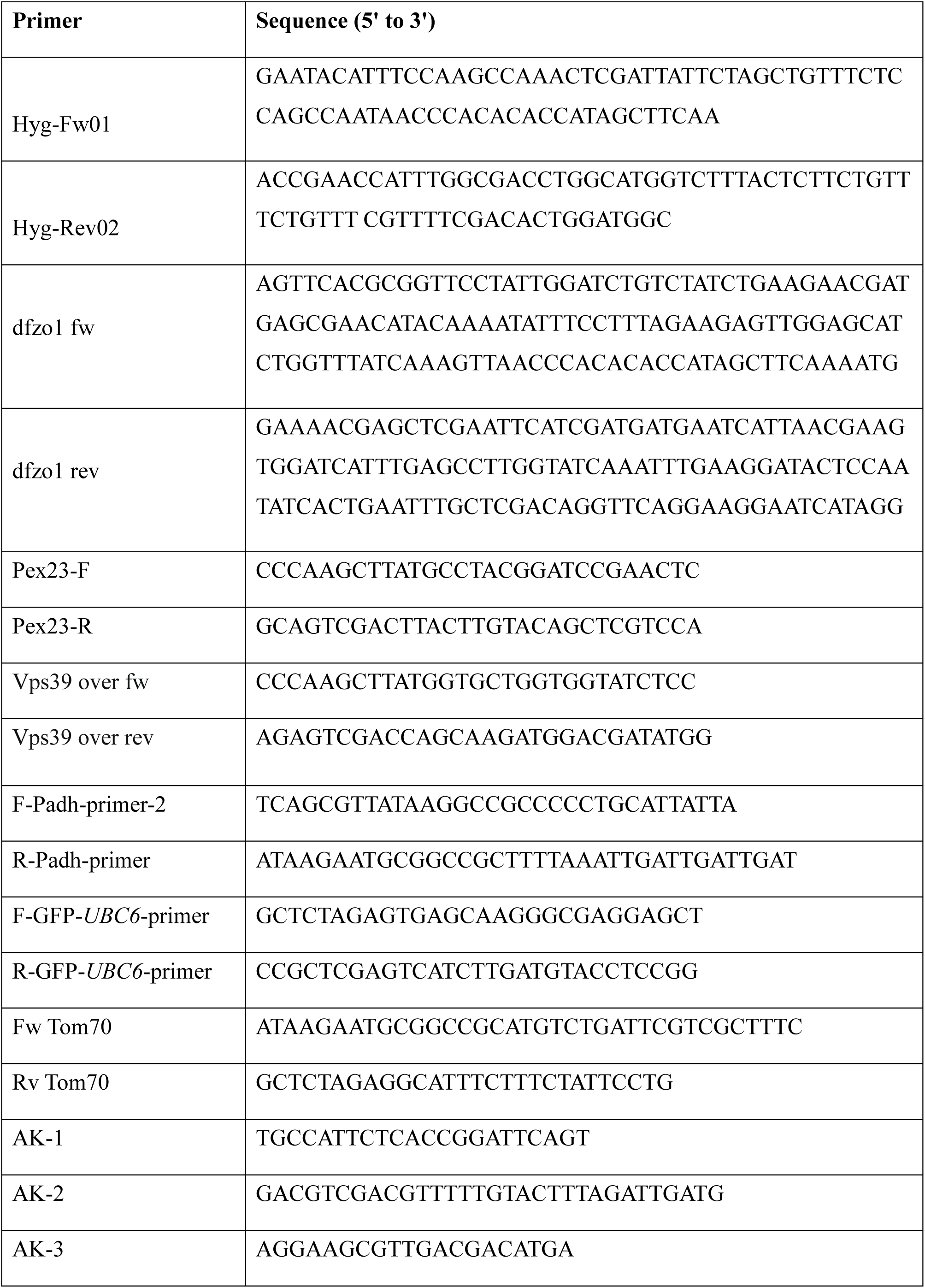
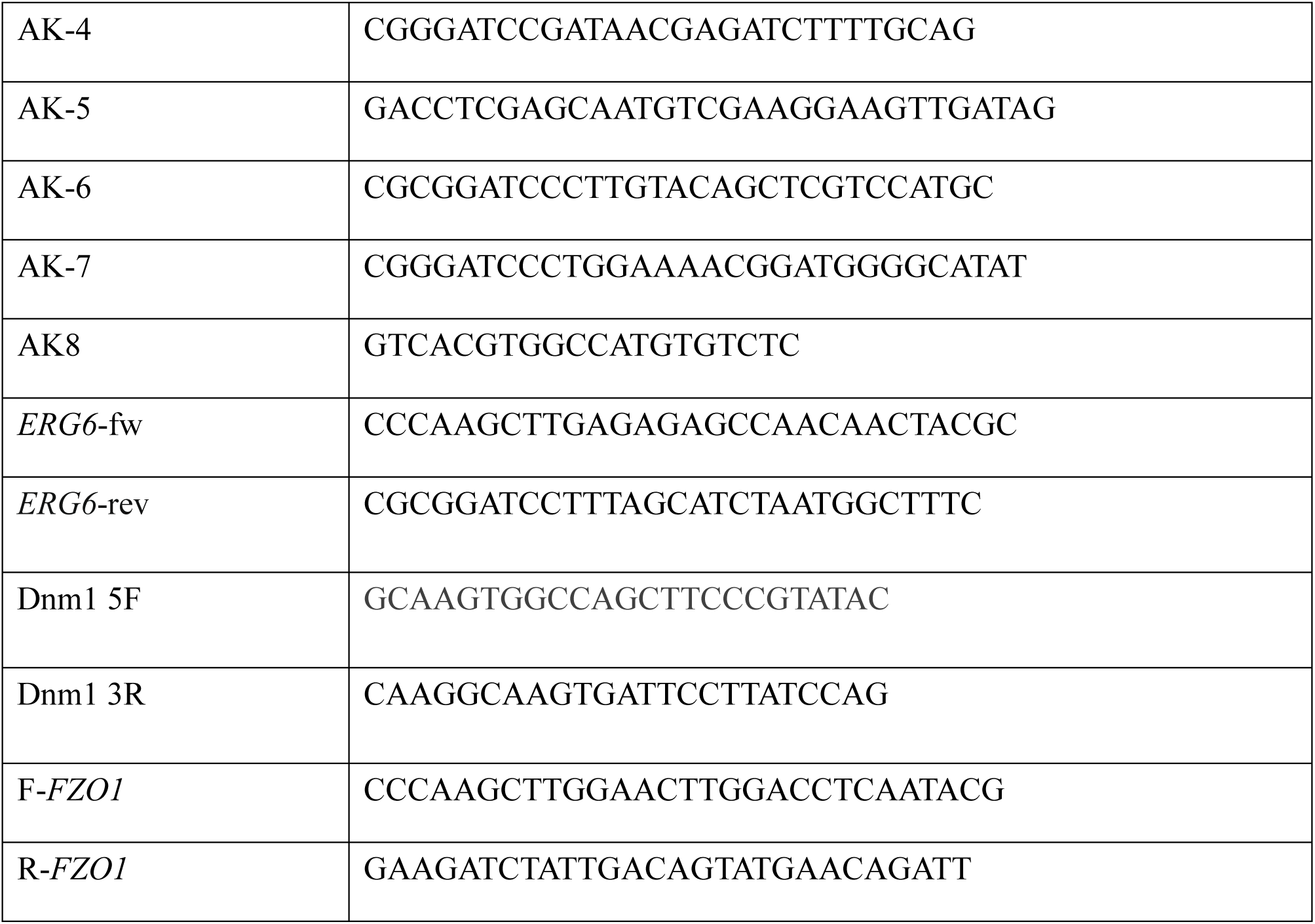
Primers used in this study.

### Construction of *fzo1* single deletion strains and *dnm1 pex23* double deletion strain

To construct an *fzo1* strain, a PCR fragment containing the *FZO1* deletion cassette was amplified with primers dfzo1 fw and dfzo1 rev using plasmid pHIPN4 as a template. The PCR product was then transformed into *yku80* cells to obtain the *fzo1* mutant.

To construct a *dnm1 pex23* double deletion strain, a PCR fragment containing the 3 kb *DNM1* deletion cassette was obtained by PCR using primers Dnm1 5F and Dnm1 3R and pDEST *DNM1-LEU* as template. The PCR product was then transformed into *H. polymorpha pex23* cells to obtain the *dnm1 pex23* double deletion strain. Correct integration was confirmed by PCR and Southern blot analysis.

### Construction of WT: P*_AOX_*Pex23-GFP::DsRed-SKL

A PCR fragment containing *PEX23*-GFP was amplified with primers Pex23-F and Pex23-R using strain Pex23-GFP as a template. The obtained PCR product was digested with *Hin*dIII and *Sal*I and inserted between the *Hin*dIII and *Sal*I sites of plasmid pHIPH4, resulting in pHIPH4-*PEX23*-GFP. *Pfl*mI-linearized pHIPH4-*PEX23*-GFP was transformed into *yku*80. *Nsi*I-linearized pHIPN4-DsRed-SKL was integrated into P*_AOX_*Pex23-GFP.

### Construction of *yku80*, *pex23* and *pex29* with an artificial ER-mitochondrion tether

To introduce an artificial ER-mitochondrion tether in the WT, *pex23* and *pex29* strains, plasmid pHIPN18 Tom70(full)-mGFP-Ubc6 was constructed. In order to get these plasmids, pHIPH18 *VPS39*, pHS6A P*_aox_*Pmp47-mGFP-Ubc6 and pHIPN Tom70 were constructed.

To construct pHIPH18 *VPS39* plasmid, a PCR fragment containing *VPS39* was amplified with primers Vps39 over fw and Vps39 over rev using genomic DNA as the template. The obtained PCR product was digested with *Hin*dIII and *Sal*I and inserted between the *Hin*dIII and *Sal*I sites of pHIPH4 plasmid, resulting in plasmid pHIPH4 *VPS39*. The *ADH1* fragment was digested with *Not*I and *Hin*dIII from pHIPN18 *PEX37* to replace the *AOX* fragment in pHIPH4 *VPS39* to obtain pHIPH18 *VPS39* plasmid.

To construct pHS6A P*_aox_*Pmp47-mGFP-Ubc6, four PCR experiments were performed. To construct plasmid pH6SA-P*_aox_*, a PCR fragment containing *AOX* promoter was amplified with primers AK-1 and AK-2 using genomic DNA as the template. The obtained PCR product was digested with *Sal*I and *Psp*XI and inserted between the *Sal*I sites of pHS6A plasmid. The *PMP47* fragment was obtained by performing PCR over the primers AK-3 and AK-4 using genomic DNA as the template. Plasmid pHIPZ-mGFP fusionator was restricted by *Hin*dIII and was performed Klenow fill in. The restricted plasmid pHIPZ-mGFP fusionator was digested by *Bgl*II, and *PMP47* fragment was digested by *Bam*HI. Two fragments were ligated to obtain plasmid pHIPZ-Pmp47-mGFP. The Pmp47mGFP fragment was amplified with primers AK-5 and AK-6 using pHIPZ-Pmp47mGFP as the template. The obtained PCR product was digested with *Psp*XI and *Bam*HI and inserted between the *Sal*I and *Bam*HI sites of plasmid pH6SA-P*_aox_*, resulting in pHS6A-P*_aox_*Pmp47-mGFP. The *UBC6* fragment was obtained by performing PCR over the primers AK-7 and AK-8 using genomic DNA as the template. The obtained PCR product was digested with *Bam*HI and inserted between the *Bam*HI and *Sma*I sites of plasmid pHS6A-P*_aox_*Pmp47-mGFP, resulting in pHS6A P*_aox_*Pmp47-mGFP-Ubc6.

To construct pHIPN Tom70, a PCR fragment contains *TOM70* was amplified with primers Fw Tom70 and Rv Tom70 using genomic DNA as the template. Use *Not*I and *Xba*I to restrict PCR fragment and plasmid pHIPN7 GFP-SKL and ligate two fragments to get pHIPN Tom70.

Plasmid pHIPN18 Tom70(full)-mGFP-Ubc6 was construct as follows. First a PCR fragment containing *ADH1* promoter was amplified with primers F-Padh-primer-2 and R-Padh-primer using plasmid pHIPH18 *VPS39* as a template. The PCR fragment was digested by enzyme *Psi*I and *Not*I, then inserted into plasmid pHIPN Tom70 to get plasmid pHIPN18-Tom70. A PCR fragment containing mGFP-*UBC6* was amplified with primers F-GFP-*UBC6*-primer and R-GFP-*UBC6*-primer using plasmid pHS6A P*_aox_*Pmp47-mGFP-Ubc6 as a template. Two *Xba*I/*Xho*I digested fragments from the obtained PCR fragment and pHIPN18-Tom70 were ligated to get plasmid pHIPN18 Tom70(full)-mGFP-Ubc6.

Then *Bst*XI-linearized pHIPN18 Tom70(full)-mGFP-Ubc6 was transformed into WT*, pex23,* and *pex29* strains to construct P*_ADH1_*Tom70-mGFP-Ubc6 (ERMIT tether) expressing strains.

### Construction of strains expressing Erg6-mGFP, Fzo1-GFP and Dnm1-GFP under control of their endogenous promoter

A plasmid encoding Erg6-GFP was constructed as follows: A PCR fragment encoding the C-terminus of *ERG6* was amplified by using primers *ERG6*-fw and *ERG6*-rev with WT genomic DNA as a template; the obtained PCR fragment was digested with *Hin*dII/*Bam*HI and ligated with the *Hin*dII/*Bgl*II digested pHIPN-Pex14-mGFP to get plasmid pHIPN-Erg6-mGFP. Subsequently, to construct the Erg6-mGFP expressing strains, *Bgl*II-linearized pHIPN-Erg6-mGFP was transformed into *yku80, pex23, pex24, pex29* and *pex32* strains.

A plasmid encoding Fzo1-GFP was constructed as follows: A PCR fragment encoding the C-terminus of *FZO1* was amplified by using primers F-*FZO1* and R-*FZO1* with WT genomic DNA as a template; Two *Hin*dII/*Bgl*II digested fragments from the obtained PCR fragment and pHIPN-Pex14-mGFP were ligated to get plasmid pHIPN-Fzo1-mGFP. Subsequently, to construct the Fzo1-mGFP expressing strains, *Bbv*CI-linearized pHIPN-Fzo1-mGFP was transformed into *yku80, pex23, pex24, pex29* and *pex32* strains.

A plasmid encoding Dnm1-GFP was constructed as follows: two *Hin*dII/*Bgl*II digested fragments from the plasmid pHIPZ-Dnm1-GFP and pHIPN-Pex14-mGFP were ligated to get plasmid pHIPN-Dnm1-mGFP. Subsequently, to construct the Dnm1-GFP expressing strains, *Bst*BI-linearized pHIPN-Dnm1-mGFP was transformed into *yku80, pex23, pex24, pex29* and *pex32* strains.

### Mitochondrial membrane potential analysis

Rhodamine 123 was used to monitor the mitochondrial membrane potential. Cells were grown on 0.5 % glucose and harvested in the log-phase and incubated with 50 nM Rhodamine123 (Invitrogen) for 30 min at 37°C. The fluorescence intensity of the cells stained with Rhodamine123 was analysed using a flow cytometer (BD Accuri™ C6 Plus) as described previously (Drakulic et al., 2005).

### Preparation of yeast TCA lysates, SDS-PAGE and Western blot analysis

Cell extracts of TCA-treated cells were prepared for SDS-PAGE as described previously (Baerends et al., 2000). Equal amounts of protein were loaded per lane and blots were probed with anti-mGFP antibodies (Santa Cruz Biotech, #sc-9996; 1:2000 dilution), anti-elongation factor 1-α (EF1A) antibodies (1:10000 dilution) (Kiel et al., 2007) or anti-pyruvate carboxylase 1 (Pyc1) antibodies (1:10,000 dilution) (Ozimek et al., 2007). Secondary goat anti-rabbit (Thermo Fisher Scientific 31460, 1:5000 dilution) or goat anti-mouse (Thermo Fisher Scientific 31430, 1:5000 dilution) antibodies conjugated to horseradish peroxidase (HRP) were used for detection. EF1A or Pyc1 were used as loading controls. Enhanced chemiluminescence (Amersham ECL Prime, #RPN2232) was used to visualize proteins of interest following the manufacturer’s instructions.

### Quantification of Western blots

Blots were scanned using a densitometer (Bio-Rad, XRS+) and protein levels were quantified using ImageJ software. The density of each band measured was standardized by dividing the density of the corresponding loading control band. Relative Density values were calculated by using the same standard sample (WT sample) as a common reference. Significance was determined using two-tailed t-test. The data shown were obtained from two/three independent experiments.

### Fluorescence microscopy

For vacuole staining, 1 ml of cell culture was supplemented with 1 μl FM4-64 (Invitrogen, #T13320), incubated for 60 min at 37°C, and analyzed. The lipid droplet dye BODIPY 493/503 (Invitrogen, #D3922) was used at a concentration of 1 µg/mL, incubated for 10 min at 37°C. Mitochondria were stained by adding 0.1 µL Mitotracker Orange (Invitrogen, #M7510; 1 mM) and Mitotracker Red (Invitrogen, #M7512; 1 mM) to 1 mL of cells. These cells were incubated for 5 min at 37 °C and subsequently spotted on agar containing glucose. DAPI (Sigma, #32670; 1 µg/mL) was used for DNA staining (de Boer and van der Klei, 2023).

Images were obtained from the cells in growth media or from 200 nm thick cryosections for CLEM using a fluorescence microscope (Axioscope A1; Carl Zeiss) using Micro-Manager software and a digital camera (Coolsnap HQ^2^; Photometrics). The GFP and BODIPY fluorescence were visualized with a 470/40 nm band-pass excitation filter, a 495 nm dichromatic mirror, and a 525/50 nm band-pass emission filter. Mitotracker and FM4-64 fluorescence were visualized with a 546/12 nm band-pass excitation filter, a 560 nm dichromatic mirror, and a 575-640 nm band-pass emission filter. DAPI fluorescence was visualized with a 380/30 nm band-pass excitation filter, a 420 nm dichromatic mirror, and a 460/50 nm band-pass emission.

Airyscan images were captured with a confocal microscope (LSM800; Carl Zeiss) equipped with a 32-channel gallium arsenide phosphide photomultiplier tube (GaAsP-PMT), Zen2009 software (Cal Zeiss) and a 63×1.40 NA objective (Carl Zeiss).

Image analysis was performed using ImageJ, and figures were prepared using Adobe Illustrator software.

### Quantification of LDs numbers

The number of LDs was quantified from Z-stacks of CLSM images. Quantification was performed automatically using a custom-made plugin from ImageJ (Thomas et al., 2015). 900 cells of three independent cultures were quantified. Significance was determined using two tailed student’s t-test.

### Electron microscopy

For morphological analysis, cells were fixed in 1.5% potassium permanganate, post-stained with 0.5% uranyl acetate and embedded in Epon. Mitochondrial area and the distance between mitochondria are measured in ImageJ.

Serial sectioning was performed on cryo-fixed cells. These cells are cryo-fixed and freeze substituted as described previously (Wu et al., 2019). 60 nm serial sections were collected on formfar coated single slot copper grids and inspected with a CM12 transmission electron microscope (TEM) (Philips).

CLEM was performed as described previously (de Boer and van der Klei, 2023). After fluorescence imaging, the grid was post-stained and embedded in a mixture of 0.5% uranyl acetate and 0.5% methylcellulose. Acquisition of the double-tilt tomography series was performed manually in a CM12 TEM (Philips). The CM12 TEM ran at 100 kV and a tilt range of 45° to -45° with 2.5° increments was included. To construct the CLEM images, pictures taken with FM and EM were aligned using the eC-CLEM plugin in Icy (Paul-Gilloteaux et al., 2017). Reconstruction of the tomograms and alignment of the serial sections was performed using the IMOD software package.

## Competing interests

The authors declare no competing or financial interests.

## Funding

This work was supported by grants from the China Scholarship Council (CSC) to H.C and F.W..

## Supplemental data

**Figure S1.**
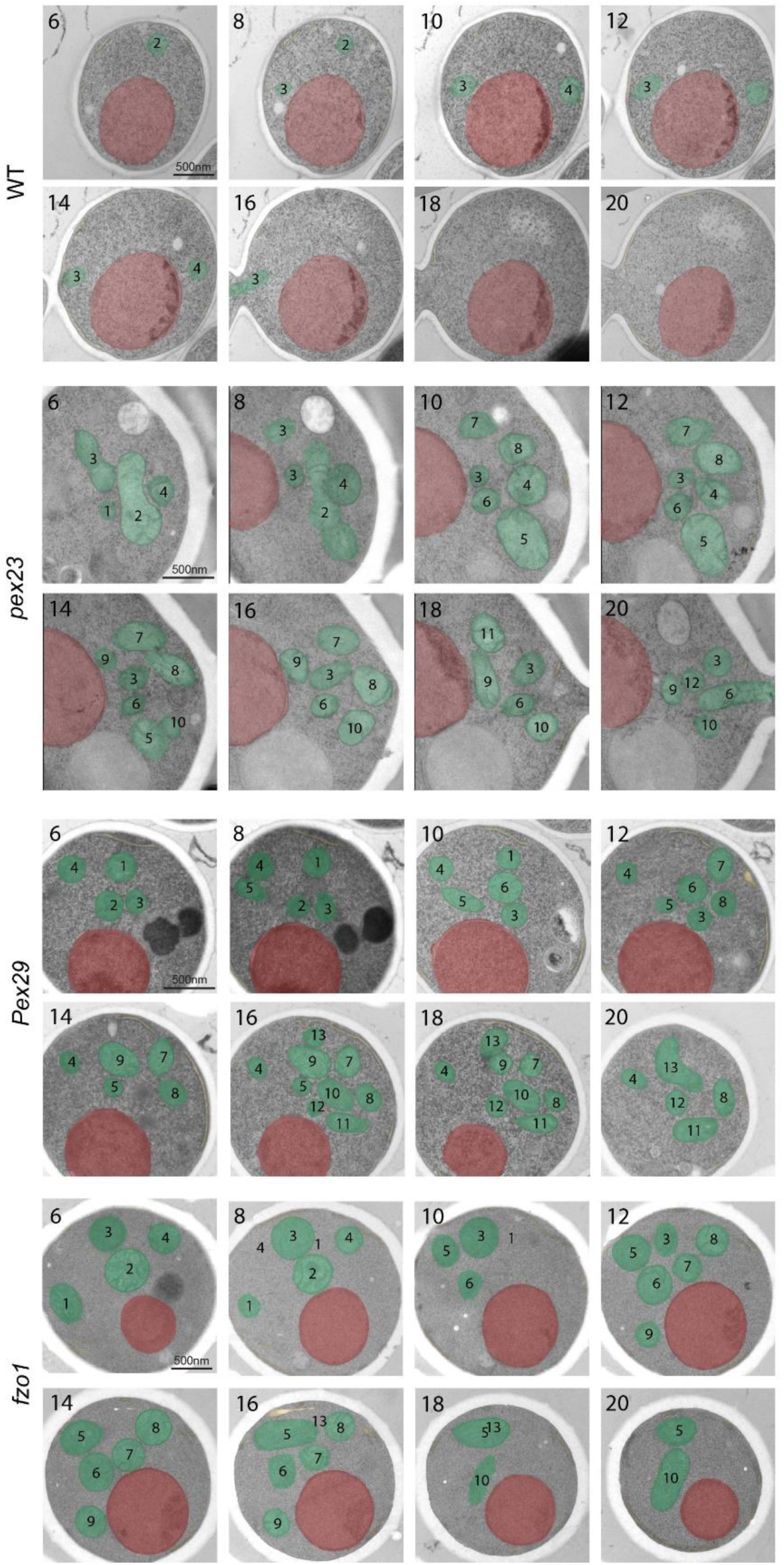
*H. polymorpha pex23, pex29* and *fzo1* cells contain a cluster of multiple mitochondria. Eight consecutive EM sections of cells grown on glucose medium. Red represents nucleus and green represents mitochondria. Numbers represent indicated mitochondria. Scale bar: 500nm.

## Notes

### Competing Interest Statement

The authors have declared no competing interest.

